# Triggering blood-lipid-profile by immunomodulatory and antimicrobial molecules for cross-evaluation of early detection of cancer

**DOI:** 10.1101/2023.02.28.530394

**Authors:** Om Prakash, Feroz Khan

**Affiliations:** Technology Dissemination and Computational Biology Division, CSIR-Central Institute of Medicinal & Aromatic Plants, P.O.-CIMAP, Kukrail Picnic Spot Road, Lucknow-226015 (Uttar Pradesh), India

**Keywords:** Cancer, Disease, Early, Prediction, Small-molecule, Stomach

## Abstract

**Background:** Al-guided mapping establishes relationship between gene-signature and phenotype. Since, Immunomodulatory & antimicrobial molecules, both show anticancer as well as their relation with lipid; therefore, can be further used for cross-validation or enrichment analysis for early detection of cancer. Considering the information, a study was performed on the influence of Immunomodulatory and antimicrobial molecules on blood lipid profile at the early stage of cancer.

**Materials and Method:** Present study established a relationship between small molecule-vs-genotype to phenotype. Genotypic features extracted through RNAseq analysis for stomach adenocarcinoma, which was further mapped with blood lipid profile. RNA-seq expression, from a population, was collected as gene expression profile of transcriptome. Gene signature was identified through in-house processing. Blood lipid profile was as: Total Cholesterol, LDL Cholesterol, HDL Cholesterol, Triglycerides, Non-HDL-C, and TG to HDL ratio.

**Results:** To evaluate the Immunomodulatory and antimicrobial molecules for anticancer activity in terms of their influence on blood lipid-profile, computational models were established for developing relationships between small molecules-vs-genotype. Immunomodulatory and antimicrobial activity showed significant impact on genotypic expression at cancerous conditions.

**Conclusion:** It was concluded that usage of Immunomodulatory small molecules & antibiotics showed their impact on blood lipid profile. These results may support cross-checking the possibility of cancer during the early detection of cancer. Furthermore, suggested the use of immunomodulatory and antimicrobial small molecules for cross evaluation of early disease prediction of different cancer types.

## INTRODUCTION

Enrichment of early detection of cancer, through clinicopathological features, can be performed. The cross-evaluated results can be further used as strong evidence during diagnosis. Diagnostic values are also considered in combination of tumor markers and blood lipids. Drug response studies on blood profile are also available for tamoxifen, endoxifen and 4-hydroxyTamoxifen. These prior studies show the evidence of changes made in blood profile due to the implementation of drug molecules. Since, at an early stage of cancer generation, blood profile is significantly affected; therefore, such changes, under the influence of molecular triggering, can be used as a cross-evaluation aspect during early diagnosis of cancer.

Lipid profile is a clinically proven parameter for evaluation of multiple cancers including colorectal, breast, ovarian and stomach etc (Kok et.al., 2011). Lipid analogs are also used as regulators in in-vitro experiments (Dong et.al., 2020). Lipids are found to be influenced with drugs. Such in-depth involvement of lipids with cancer, suggests its utility for early diagnosis of cancer. Impact of hormones on lipid level is known through oestrogen. Hormonal treatment of breast cancer reduces the level of blood profile (Bundred et.al., 2005). Therefore, blood profile along with percentage of dyslipidemia are observed before chemotherapy (Xu et.al., 2020). Research is going on to understand the association between blood lipid level and cancer. Such studies consider variation in blood lipid profile including ‘total cholesterol’, triglycerides, and HDL/LDL level (Yang et.al., 2020). Blood lipid level is also observed along with other clinicopathological parameters as blood glucose and extent of inflammation (Li et.al., 2019). All these studies suggest that abnormal blood lipid profiles contain association with occurrence of cancer (Halton et.al, 1998). Blood lipid profiles are regularly estimated, if a cancer patient is under endocrine therapies (Engan T, 1996). Other studies like phospholipid level & peroxidation estimation are known to understand the malignant neoplasma as in case of breast & uterine cancer (Kotrikadze et.al., 1987). Drug response studies on blood profile are also available for tamoxifen, endoxifen and 4-hydroxyTamoxifen (Siqueria et.al., 2021). In totality, It can be said that potential correlation exists between blood lipid and cancer (Jin et.al., 2020). Diagnostic values of blood lipid profile are also considerable in combination of tumor markers and blood lipid (Jiang et.al., 2021). These studies prepare a ground for establishing a relationship between genotype (tumor markers) and phenotype (blood lipid profile) for observation of import of variation in blood lipid and the cancer status.

RNA expressions are the key material for research in several areas of clinical diagnosis (Dube et.al., 2019), including cancer (Lorenzi et.al., 2019). Categories of RNA as: miRNA (Liu et.al., 2021), circRNA expression (Dube et.al., 2019), long noncoding RNA expression (Lorenzi et.al., 2019), mRNA expression (Ruan et.al., 2020) are being used in diagnosis. Similar to DNA & protein biomarkers, RNA biomarkers are also of potential use to discriminate diseased-vs-normal conditions (Wang et.al., 2020). Therefore, transcriptome sequencing is used in diagnosis of various disorders (Akula et.al., 2021). Transcriptome RNA expressions are also used for identification of targetable disease regulatory networks for disease diagnosis (Gaffo et.al., 2021). RNA based studies are well known with hepatocellular carcinoma (Chen et.al., 2018), colorectal cancer (Mamelli et.al., 2021), pathway detection (Wu et.al., 2021), prostate tumor (Xia et.al., 2018), and breast cancer (Liu et.al., 2021) etc. RNA expression profile/signature are also utilised for early diagnosis (Zheng et.al., 2019; Liu et.al., 2021). Therefore, it is obvious that, Early diagnosis is directly linked with development of biomarker signature (i.e., gene set) containing population discrimination capacity (Yang et.al., 2020). Such studies provide clinically relevant results for diagnosis (Brakenridge et.al., 2021). These studies provide a ground for accessing RNA expressions as genotypic features for early detection of cancer. Genotype-to-phenotype relation is used also for linking lipid metabolism with cancer related genes (Kale et.al., 2020).

Stomach adenocarcinoma (STAD) is one of the leading causes of deaths in the world. Lack of early detection of STAD is also a major barrier behind present status. Biomarker genes, linked with extracellular matrix and platelet-derived growth factor, are suggested in early detection of STAD (Tan et.al., 2021). Besides this, CDCA7-regulated inflammatory mechanism (Guo et.al., 2021), angiogenesis-related lncRNAs (Han et.al., 2021), PLXNC1 an immune-related gene (Ni et.al., 2021), and Ferroptosis-related gene (Xiao et.al. 2022) etc. are also claimed for early detection of STAD. Different stages of STAD are known through prognostic models from the TCGA database (Qian et.al., 2022). Various combinations of gene-signature genes still remain to explore, which creates an opportunity to reveal aspects for early detection of STAD.

In present study, a link between blood lipid profile (as phenotype) and gene signature (as genotype) derived for stomach adenocarcinoma, used to predict the possible changes/ status of cancer. Established relationship between genotype and phenotype, helped us on estimation of blood-lipid-profile in reference of gene-signatures. Since, Immunomodulatory & antimicrobial molecules, both show anticancer as well as their relation with lipid; therefore, can be further used for cross-validation or enrichment analysis for early detection of cancer. Considering the information, we performed a study on the influence of Immunomodulatory and antimicrobial molecules on blood lipid profile at the early stage of cancer. Inference extracted from this process can be directly used during early diagnosis of stomach adenocarcinoma.

## MATERIALS AND METHOD

Implementation of triggering-impact of small molecules on blood profile, was performed by establishment of AI-guided relation between blood lipid-profile (clinicopathological phenotype) and expression of gene signature. Blood lipid profile has already been proven for their performance on discrimination between cancer and normal status of the cell (doi: https://doi.org/10.1101/2023.02.14.23285899). Workflow behind the protocol is (Figure 1):

**Figure 1.**
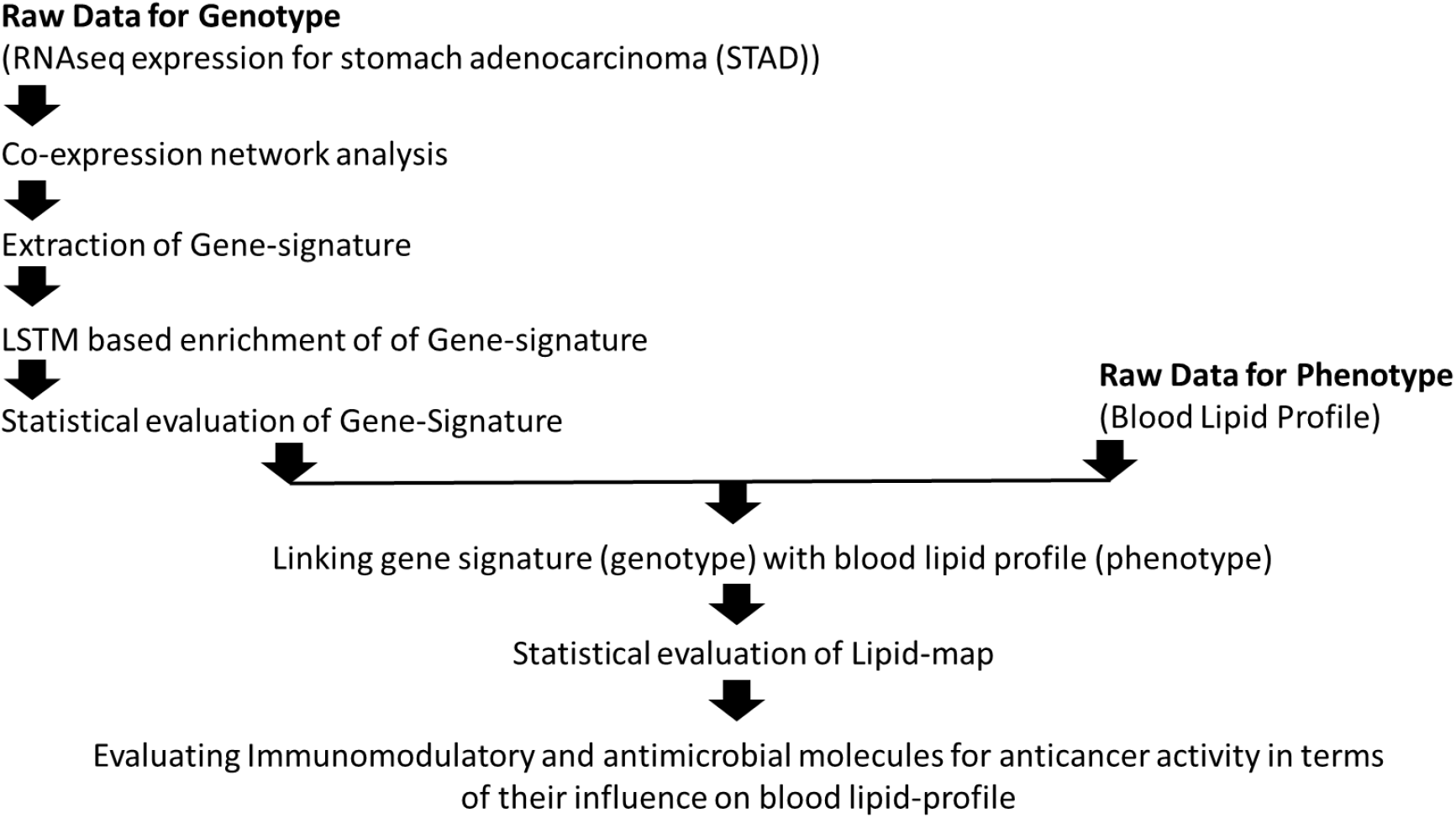
Workflow for assessment of Immunomodulatory and antimicrobial molecules for anticancer activity in terms of their influence on blood lipid-profile as clinicopathological feature

### Genotype & phenotype features and their mapping

Detailed description of methodology for extraction of genotype features can be accessed from previous study (doi: https://doi.org/10.1101/2023.02.14.23285899). Gene signature must bear discrimination capacity between diseased and non-diseased population. In brief, Raw data was collected as RNAseq expression for stomach adenocarcinoma from ‘The Cancer Genome Atlas’ (TCGA) derived Gene Expression Profiling Interactive Analysis database. Population sample size was: 408 independent tumor samples and 211 matching normal tissue samples. DEGs mapped with plasmaproteome databased (pDEGs) used for analysis. DEGs identified at ‘Fold change (log_2_)’ of 1.5 and ‘Significance (-log_10_)’ boundary at 0.05. pDEGs processed for co-expression network analysis (Vizeacoumar et.al., 2021). Network analyjsis resulted into similarity and dissimilarity network maps. pDEGs present in any of these networks can be considered as co-expressed genes; and therefore, will be assumed to be involved in existence of system model representing state of disease. Here similarity matrix was used for further analysis. Further enrichment of co-expression network was performed through simulation of systems model defined by co-expressed network map. System model simulation performed with Long Short Term Memory Recurrent Neural Network (LSTM-RNN) model (implemented in R). Identified gene signature as well as enriched gene signature were evaluated on the ground of population discrimination capacity through survival plot. Logrank p-value < 0.05 was considered as threshold for signaificant gene signature. Beyond genotype features, phenotypic features, Blood-profile was considered as phenotype for the study. General blood lipid profile used in this study, as: Total Cholestrol: 200-239(mg/dL); LDL Cholestrol: 130-159 (mg/dL); HDL Cholestrol: 40-60 (mg/dL); Triglycerides: 150-199 (mg/dL); Non-HDL-C: 130-159 (mg/dL); and TG to HDL ratio: 3.0-3.8 (mg/dL). Mapping of gene-signature and blood-lipid-profile through 740 datapoints generated. Since gene-expression was pre-classified, therefore data discrimination capacity was transformed into phenotype through data mapping for further generating classified mapped lipid-profile. Finally genotypic & phenotypic dataset sets used for development of map-model with framework of feed-forward backpropagation Artificial Neural Network (FFBP-ANN). Inputs of 06 nodes from lipid-profiles mapped with outputs of 5 nodes from enriched pDEGs. At the end, the mapped lipid-data points were evaluated for its discrimination capacity through two class multiple perceptron classification model evaluation. Statistical analysis performed to evaluate the classification model through ROC plot and AUC calculation.

### Establishing relationship between impacts of small molecule on Genotype-to-phenotype

Establishing relationship between small molecules with genotype, considered that small molecule binds to its target to perform % inhibition. Since the target protein expression are also linked with co-expressed genes, therefore co-expressed genes also considered to be modulated in the similar passion as target. In present study, immuno-modulators as well as antibiotics are considered as small molecules impacting on genotype. The modulated gene-expression, by immuno-modulators as well as antibiotics, further passed through Global genotype-vs-phenotype AI-model to obtain the relative systemic impact on the phenotypic blood-lipid profile.

## RESULTS AND DISCUSSION

### Gene-signature, identified through network analysis and AI-guided simulation of systems model

To identify gene-signature, firstly DEGs expressed in blood plasma were picked (Tabulated in supplementary Table-1). Total 39 plasma-DEGs collected. Next step was identification of co-expressed pDEGs. Network analysis method used for identification of co-expressed gene-set based on similarity and dis-similarity matrix generated through network analysis. Similarity matrix-based network contain 11-genes (out of 39, AFF3, APBB1, C5, CHRD, COL4A5, EEF1A2, ZBTB16, IL1RL1, GFRA3, ZNF662, and MAGED4B); while dissimilarity matrix-based network contains 10 genes (out of 39, AFF3, C5, CHAF1A, EFNA3, SLC52A3, LOX, ILDR1, GFRA3, STC1, and ZNF662). Both networks bear 4 genes (AFF3, C5, GFRA3 and ZNF662) common. Similarity matrix-based network was the first significant signature with population discrimination capacity of p-value 6E-05. It was further evaluated for enriched filtering of the network through the LSTM system model. Similarity network used for development of LSTM system model. LSTM model simulation, and found only 05 genes to be involved in simulation. The AFF3 gene is known to be a putative transcription activator that may function in lymphoid development and oncogenesis. APBB1 gene is a Amyloid beta precursor, its role is in response to DNA damage & apoptosis. C5 gene is a Pyrimidine/ purine nucleoside phosphorylase. CHRD gene is a key development protein, and is used during early embryonic tissue and also expression cancer conditions. The COL4A5 gene is collagen alpha, it is used as constituent of extracellular matrix. Reactome based pathway enrichment analysis showed that signatures of genes belong to the ‘Extracellular Matrix Organization’, ‘immune system’, ‘DNA repair’ and ‘developmental biology’. It is shown that the most significant impact on stomach adenocarcinoma is due to genes involved in extracellular matrix organization. In a prior study, STAD early detection biomarker genes were suggested from extracellular matrix [30](Tan et.al., 2021). These observations are also in compliance with prior studies on development of gastric cancer (Moreira et.al., 2021). These 05 genes were further used for establishing relation with blood profile.

Population discrimination capacity of signature was evaluated with survival plot. Two gene signatures were evaluated. First with 11 gene-set and second with 05 gene-set. Second one was filtered from the first one. Filtering was performed with LSTMl based simulation of the system model. First gene signature showed logrank p-value of 6e-05. while the Second one also gained the significant p-value of 0.0046; both were less than 0.05 (doi: https://doi.org/10.1101/2023.02.14.23285899). To link gene signatures with blood profile, gene expressions were mapped into 6 parameters from blood lipid profile. Parameters were: Total Cholesterol: 200-239(mg/dL); LDL Cholesterol: 130-159 (mg/dL); HDL Cholesterol: 40-60 (mg/dL); Triglycerides: 150-199 (mg/dL); Non-HDL-C: 130-159 (mg/dL); and TG to HDL ratio: 3.0-3.8 (mg/dL). Since the mapped data must contain population discrimination capacity therefore to evaluate performance of a model an ANN based classifier was Trained and evaluated. Architecture of ANM multiple perceptron model was 6-4-2 with learning rate of 0.3 and momentum of 0.2 for 500 epochs. Developed model performed with accuracy of more than 99% (AUC-0.99). Furthermore, the model was also cross validated with 10-fold cross validation, with gain of 99.46% of accuracy during classification of 740 instances (Supplementary Table 2, 3 & 4).

### Estimating impact of immunomodulatory small molecule on Genotype, followed by phenotype

Prior study results indicated that, there is a significant fall in density of lipid in blood in cancerous conditions than normal. Therefore, this result is used as a general rule for early detection of cancer. But, lipid-density may also go down for many physiological reasons, then how can we confirm that that variation in blood-lipid profile is linked with any existence of cancer? Therefore, to cross-validate, external impulses, from immunomodulatory and antimicrobial small molecules (antibiotics), used to trigger the genotype. Phenotypic impact observed through genotype-phenotype mapped model. Small molecule modulated genotype expressions were passed through the global-genotype-vs-phenotype model, and phenotypic influence found. Modulation based on %inhibition of any small molecule showing prime target to COX2. It should be noticed that here no description about dosage was given, which may be a part of further research. Here, COX2 (PTGS2) co-express with IL1B, CXCL1, CXCL1 and CXCL8. Since in our dataset only CXCL1 & CXCL8 was expressed, therefore COX2 was considered as the function of CXCL1 & CXCL8. Considering the expression in cancer condition, expression of both co-expressed genes is reduced by 50% (CXC1: 50.086 -> 25.00; CXCL8: 30.630 -> 15.00) before further simulation. As it was seen that, immunomodulatory activity through co-expressed gene-set, in normal condition, was higher than tumor condition. After considering the impact of immunomodulatory small molecules by inhibition of 50%, expression of the gene-set was found to be approaching the normal condition (Table 1).

**Table 1.** Impact of immunomodulatory (targeting COX2) small molecules on phenotype. Tumor condition expression is shown in reference of gene-set for COX2 co-expressed genes.

| COX2 co-expressed genes | "PTGS2", "IL1B", "CXCL2", "CXCL1", "CXCL8" |  |  |  |  |  |
| --- | --- | --- | --- | --- | --- | --- |
| Non-tumor condition | 214.2378 | 174.1326 | 47.65243 | 270.8775 | 169.9518 | 3.69331 |
| Tumor (before treatment) | 192.1202 | 120.379 | 35.0779 | 130.674 | 118.1375 | 2.620049 |
| Tumor after treatment with immunomodulatory | 195.0677 | 124.6875 | 37.53092 | 133.8285 | 123.6695 | 2.735683 |

### Estimating impact of antimicrobial small molecule on Genotype, followed by phenotype

Modulated genotype expression passed through Global-genotype-vs-phenotype model, and observed the phenotypic influence. Modulation, based on %inhibition of any small molecule, showed prime target to TOP2B. It should be noticed that no description about dosage was given, which may be a part of further research. Here, TOP2B co-expressed with SMC4, therefore TOP2B was considered as the function of SMC4. Considering the expression in cancer condition, expression of SMC4 reduced by 50% (SMC4: 40.415 -> 20.00) before further simulation. Phenotypic expression of co-expressed gene-set for anticancer activity in normal condition was higher than tumour condition. After considering the impact of anticancer small molecule by inhibition of 50%, gene-set influenced expression found to be approaching the normal condition (Table 2).

**Table 2.**
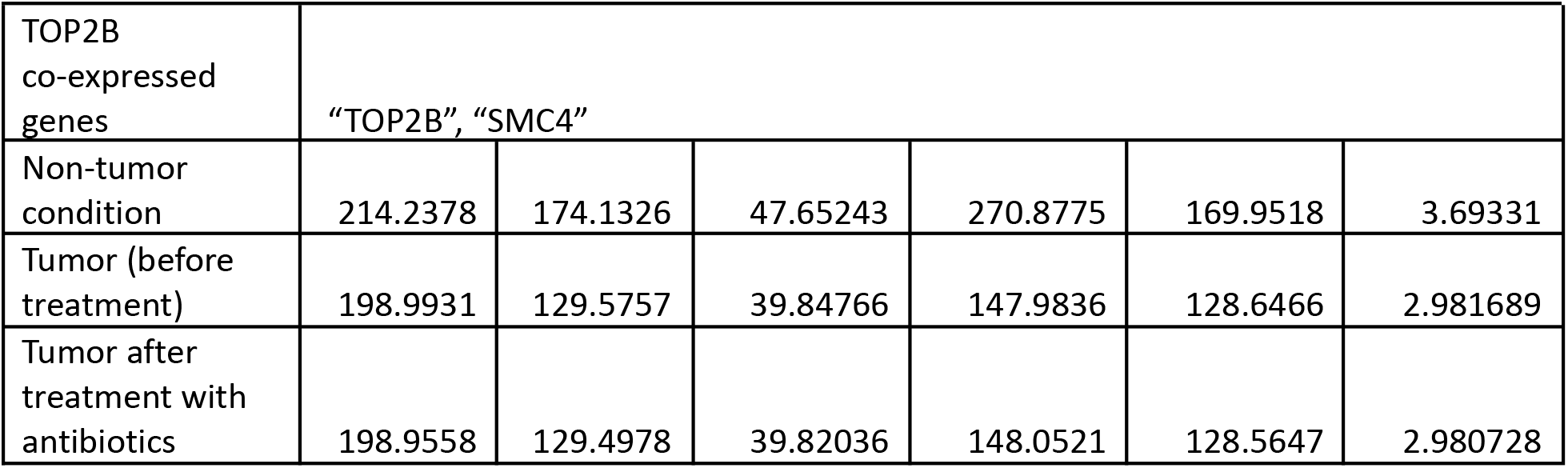
Estimating impact of antibiotics (targeting TOP2B) small molecules on phenotype. Tumour condition expression is shown in reference of gene-set for TOP2B co-expressed genes.

| TOP2B co-expressed genes | "TOP2B", "SMC4" |  |  |  |  |  |
| --- | --- | --- | --- | --- | --- | --- |
| Non-tumor condition | 214.2378 | 174.1326 | 47.65243 | 270.8775 | 169.9518 | 3.69331 |
| Tumor (before treatment) | 198.9931 | 129.5757 | 39.84766 | 147.9836 | 128.6466 | 2.981689 |
| Tumor after treatment with antibiotics | 198.9558 | 129.4978 | 39.82036 | 148.0521 | 128.5647 | 2.980728 |

Immunomodulatory molecules and antibiotics showed their impact on blood lipid profile. Both showed shifting of expression from cancerous to normal direction, i.e., reorganisation of cell signalling towards normal. These results may support cross-checking the possibility of cancer during the early detection of cancer. Furthermore, suggested small molecules can be used for early disease prediction of stomach adenocarcinoma as we have different cancer types.

## Conclusions

In present study, impact of small molecules was cross-evaluated for early detection of cancer by showing their triggering impact on blood lipid-profile. Here the impact of Immunomodulatory and antimicrobial small molecules used in terms of their influence on blood lipid-profile. AI-guided computational models take inputs of impact small molecules and click genotypes to express variation in phenotype. The identified results will support cross-evaluation of the possibility of cancer during the early detection of STAD and for different cancer types.

## Acknowledgement

Authors are thankful to the Director, CSIR-Central Institute of Medicinal & Aromatic Plants (CIMAP), Lucknow, India for infrastructure & research facilities support. Author OP is thankful to the Indian Council of Medical Research (ICMR), New Delhi, India for financial support through RA fellowship (Award letter no. BMI/11(12)/2020, dated: 04/02/2021). Author OP is also thankful to Prof. Thiol Gross, Helmholtz Institute for Functional Marine Biodiversity (HIFMB) at University of OLDENBURG, Germany and Dr. Amit Singh, Chennai Mathematical Institute (CMI) Chennai, India for fruitful suggestions on evaluation of stability of time series systems model. The CSIR-CIMAP publication number of this manuscript is CIMAP/PUB/ 2023/30.

---

**Supplementary Table 1.** Plasma-expressed gene set for signature extraction. Differentially expressed genes were mapped along plasma proteome. 39 genes, out of 62, were found to be expressed in blood plasma. Considering the feasibility of patient sampling through blood, plasma proteome expressions were strategically used for signature development. Here in the table each gene symbol is marked with ‘node ID’ from 1 to 39. (doi: https://doi.org/10.1101/2023.02.14.23285899)

| Node ID<br>-vs-Gen<br>e | PPD ID | Gene symbol | Gene name | Entrez<br>gene ID |
| --- | --- | --- | --- | --- |
| 1 | HPRD_03272 | AFF3 | AF4/FMR2 family, member 3 | 3899 |
| 2 | HPRD_04087 | APBB1 | amyloid beta (A4) precursor protein-binding, family B, member 1 (Fe65) | 322 |
| 3 | HPRD_00134 | APOD | apolipoprotein D | 347 |
| 4 | HPRD_00405 | C5 | complement component 5 | 727 |
| 5 | HPRD_03148 | CHAF1A | chromatin assembly factor 1, subunit A (p150) | 10036 |
| 6 | HPRD_04592 | CHRD | chordin | 8646 |
| 7 | HPRD_02363 | COL4A5 | collagen, type IV, alpha 5 | 1287 |
| 8 | HPRD_06012 | DPP3 | dipeptidyl-peptidase 3 | 10072 |
| 9 | HPRD_00510 | DPT | dermatopontin | 1805 |
| 10 | HPRD_04265 | EEF1A2 | eukaryotic translation elongation factor 1 alpha 2 | 1917 |
| 11 | HPRD_03225 | EFNA3 | ephrin-A3 | 1944 |
| 12 | HPRD_03957 | PDK4 | pyruvate dehydrogenase kinase, isozyme 4 | 5166 |
| 13 | HPRD_06685 | PRSS3 | protease, serine, 3 | 5646 |
| 14 | HPRD_00340 | VCAN | versican | 1462 |
| 15 | HPRD_03642 | NRP1 | neuropilin 1 | 8829 |
| 16 | HPRD_06714 | SEC23A | Sec23 homolog A (S. cerevisiae) | 10484 |
| 17 | HPRD_09843 | SLC52A3 | solute carrier family 52, riboflavin transporter, member 3 | 113278 |
| 18 | HPRD_01418 | SERPINE1 | serpin peptidase inhibitor, clade E (nexin, plasminogen activator inhibitor type 1), member 1 | 5054 |
| 19 | HPRD_13296 | FAM20A | family with sequence similarity 20, member A | 54757 |
| 20 | HPRD_11762 | ZBTB16 | zinc finger and BTB domain containing 16 | 7704 |
| 21 | HPRD_01475 | PTPN6 | protein tyrosine phosphatase, non-receptor type 6 | 5777 |
| 22 | HPRD_01087 | LOX | lysyl oxidase | 4015 |
| 23 | HPRD_03123 | IL1RL1 | interleukin 1 receptor-like 1 | 9173 |
| 24 | HPRD_09968 | G0S2 | G0/G1switch 2 | 50486 |
| 25 | HPRD_09545 | NREP | neuronal regeneration related protein | 9315 |
| 26 | HPRD_00552 | NT5E | 5'-nucleotidase, ecto (CD73) | 4907 |
| 27 | HPRD_05081 | GPNMB | glycoprotein (transmembrane) nmb | 10457 |
| 28 | HPRD_01903 | ITGAV | integrin, alpha V | 3685 |
| 29 | HPRD_13735 | ILDR1 | immunoglobulin-like domain containing receptor 1 | 286676 |
| 30 | HPRD_10420 | GFRA3 | GDNF family receptor alpha 3 | 2676 |
| 31 | HPRD_03115 | STC1 | stanniocalcin 1 | 6781 |
| 32 | HPRD_04183 | IGFBP7 | insulin-like growth factor binding protein 7 | 3490 |
| 33 | HPRD_16291 | PPP1R14A | protein phosphatase 1, regulatory (inhibitor) subunit 14A | 94274 |
| 34 | HPRD_02698 | FABP4 | fatty acid binding protein 4, adipocyte | 2167 |
| 35 | HPRD_01981 | PCCA | propionyl CoA carboxylase, alpha polypeptide | 5095 |
| 36 | HPRD_03774 | PER1 | period circadian clock 1 | 5187 |
| 37 | HPRD_13516 | ZNF662 | zinc finger protein 662 | 389114 |
| 38 | HPRD_06631 | MAGED4B | melanoma antigen family D, 4B | 81557 |
| 39 | HPRD_11750 | GPX3 | glutathione peroxidase 3 (plasma) | 2878 |

**Supplementary Table 2.** Statistics showing evaluation through 10-fold cross validation of two-class classification model for evaluation of discrimination capacity of lipid-profile

|  | TP | FP | Precision | Recall | F-Measure | MCC | ROC area | PRC area | Class |
| --- | --- | --- | --- | --- | --- | --- | --- | --- | --- |
|  | 0.997 | 0.008 | 0.992 | 0.997 | 0.995 | 0.989 | 1 | 1 | T |
|  | 0.992 | 0.003 | 0.997 | 0.992 | 0.995 | 0.989 | 1 | 1 | N |
| Weighted Avg. | 0.995 | 0.005 | 0.995 | 0.995 | 0.995 | 0.989 | 1 | 1 |  |
| Correctly Classified Instances 736 99.4595 % |  |  |  |  |  |  |  |  |  |
| Incorrectly Classified Instances 4 0.5405 % |  |  |  |  |  |  |  |  |  |
| Kappa statistic 0.9892 |  |  |  |  |  |  |  |  |  |
| Mean absolute error 0.0091 |  |  |  |  |  |  |  |  |  |
| Root mean squared error 0.0631 |  |  |  |  |  |  |  |  |  |
| Relative absolute error 1.8151 % |  |  |  |  |  |  |  |  |  |
| Root relative squared error 12.6272 % |  |  |  |  |  |  |  |  |  |
| Total Number of Instances 740 |  |  |  |  |  |  |  |  |  |

**Supplementary Table 3.** Phenotypic expression related with Normal cell

| Sig1 | Sig2 | Sig3 | Sig4 | Avg (Normal Signature) |
| --- | --- | --- | --- | --- |
| 214.2378 | 235.3203 | 346.6813 | 219.0083 | 253.8119 |
| 174.1326 | 146.494 | 169.8594 | 142.7449 | 158.3077 |
| 47.65243 | 49.63761 | 58.09198 | 57.39752 | 53.19489 |
| 270.8775 | 186.109 | 251.5707 | 170.4471 | 219.7511 |
| 169.9518 | 147.512 | 241.0234 | 156.4613 | 178.7371 |
| 3.69331 | 4.04113 | 6.303563 | 3.182708 | 4.305178 |

**Supplementary Table 4.** Phenotypic expression related with Tumor cell

| Sig1 | Sig2 | Sig3 | Sig4 | Avg (Tumor Signature) |
| --- | --- | --- | --- | --- |
| 207.9712 | 221.4865 | 279.8497 | 215.6043 | 231.2279 |
| 170.757 | 143.7998 | 157.0263 | 144.6829 | 154.0665 |
| 45.91087 | 46.73759 | 50.49587 | 45.25915 | 47.10087 |
| 227.1245 | 187.4013 | 202.5042 | 172.4644 | 197.3736 |
| 149.5762 | 150.1777 | 196.4386 | 144.9133 | 160.2765 |
| 3.34872 | 3.914579 | 4.701655 | 3.213511 | 3.794616 |

